# Acute and 90-Day Repeated-Dose Oral Toxicity of *Urtica dioica* L. Leaf Extract Granules in Rodents

**DOI:** 10.64898/2026.10.20.760560

**Authors:** Nitya Nurul Fadilah, Srie Rezeki Nur Endah, Sri Adi Sumiwi, Widya Nurul Aini, Muhamad Ilham

## Abstract

*Urtica dioica* L. leaves are traditionally used in Indonesia for gout-related complaints, and granules formulated from their ethanolic leaf extract have shown xanthine oxidase inhibitory and uric acid-lowering activity; the safety of the finished formulation, however, remains uncharacterised. This study evaluated acute and repeated-dose oral toxicity under Indonesian National Agency of Drug and Food Control (BPOM) Regulation No. 10 of 2022. Female Swiss Webster mice received a single dose of 5, 50, 500, or 5000 mg/kg granules, and Sprague-Dawley rats of both sexes (five per sex per group) received 77.5, 155, or 310 mg/kg/day for 13 weeks (90 days), with satellite groups observed through a 30-day recovery period. No mortality occurred at 5000 mg/kg, giving an LD₅₀ above that limit; clinical signs, body weight, and rectal temperature showed no dose-related change. After repeated dosing, erythrocyte indices, blood urea nitrogen, and creatinine were unchanged in both sexes. Leukocyte count rose in high-dose females (p < 0.05) but was not observed in the high-dose recovery group, and liver weight differed among female groups without a dose-response pattern. Weight gain at the highest dose differed from control in both sexes but in opposite directions. Control values outside the strain reference range and baseline body-weight imbalance preclude establishing a no-observed-adverse-effect level.

## Introduction

Hyperuricemia affects a growing share of adults worldwide and precedes gout, urolithiasis, and chronic kidney disease [1]. Pharmacological control relies largely on xanthine oxidase inhibitors, of which allopurinol remains the most widely prescribed, but its use is constrained by hypersensitivity reactions, dose adjustment in renal impairment, and variable adherence during long-term therapy [2–4]. These limitations have sustained a large market for plant-based preparations marketed for uric acid control. Such products are typically consumed daily over months, without medical supervision and without the dose standardisation applied to registered medicines, so that the duration and magnitude of exposure may exceed anything evaluated in the pharmacological studies used to justify their use [5,6].

*Urtica dioica* L. (Urticaceae), known in Indonesia as Jelatang, is among the species used traditionally for gout-related complaints, with leaf decoctions taken to relieve joint pain and swelling. Phytochemical surveys of the leaves report flavonoids, phenolic acids, tannins, saponins, and triterpenoids [7], and flavonoids in particular are established inhibitors of xanthine oxidase, the enzyme catalysing the final two steps of purine catabolism[8,9]. Extracts of the species have shown uric acid-lowering activity in rodent models of induced hyperuricemia, providing a mechanistic basis for the traditional indication rather than merely an ethnobotanical one [6].

Safety data for the genus remain limited relative to this pharmacological interest. Published toxicity assessments of *Urtica* species have largely examined crude aqueous or hydroalcoholic extracts after single administration or short-term exposure of up to 28 days [10],[11], and have generally reported no mortality at doses of 2000 mg/kg or above. Such designs establish an acute margin of safety but do not address the questions that matter for a product intended for prolonged use: whether repeated daily exposure produces cumulative haematological or hepatorenal change, and whether any change that does appear resolves once dosing stops. Delayed effects are of particular concern for plant preparations rich in polyphenols, since these constituents and their metabolites can accumulate in liver and kidney tissue [12].

Development of *U. dioica* towards a standardised dosage form has meanwhile advanced. We previously formulated the ethanolic leaf extract into oral granules and screened several compositions for xanthine oxidase inhibition; the formulation containing 5% (w/w) extract gave the strongest inhibition, with an IC_50_ of 47.18 µg/mL, and significantly lowered serum uric acid in potassium oxonate-induced mice [13]. That work established pharmacological proof of concept and a candidate effective dose, and it defined the specific material, a granule rather than a crude extract, that any subsequent safety assessment would need to address.

The toxicological profile of this formulation has not been established, and it cannot be established solely from studies of crude *Urtica* extracts [14]. Granulation alters the physical form in which the extract reaches the gastrointestinal tract and introduces excipients, including lactose, polyvinylpyrrolidone, and maltodextrin, that are absent from the plant material and that may modify dissolution and absorption. A safety assessment of the finished product is therefore a prerequisite for further development [15].

This study evaluated the acute oral toxicity of *U. dioica* leaf extract granules in mice and their repeated-dose oral toxicity over 13 weeks in rats of both sexes, following the Indonesian regulatory framework for preclinical toxicity testing [9]. Doses for the repeated-dose phase were derived from the pharmacologically effective dose established in our earlier work, and satellite groups were maintained through a 30-day recovery period to assess the reversibility of any change observed and to detect effects emerging after treatment ceased. The aim was to define the exposure conditions under which the formulation can be considered safe for prolonged use, and to identify the endpoints requiring further study before human exposure is contemplated [16].

## Materials and Methods

### Plant Material and Authentication

Leaves of *Urtica dioica* L. (Urticaceae) were collected in January 2026 from Sukasetia Village, Cisayong District, Tasikmalaya Regency, West Java, Indonesia (−7.240423, 108.128044). Botanical identity was authenticated by the Plant Taxonomy Laboratory, Department of Biology, Faculty of Mathematics and Natural Sciences, Universitas Padjadjaran (determination certificate No. 19/HB/01/2026).

### Extraction

Fresh leaves (5 kg) were air-dried to yield 625 g of dried material. A 500 g portion was milled and passed through a 40-mesh sieve, then extracted with 96% ethanol at a drug-to-solvent ratio of 1:10 by remaceration over three successive 24-h cycles. The pooled filtrate was concentrated under reduced pressure at 50 °C, giving a viscous extract with a yield of 16.25 % (w/w of dried material), which was stored in a tightly closed container at 25 °C until use [17].

### Phytochemical Characterization

Qualitative screening for alkaloids, flavonoids, saponins, tannins, quinones, steroids, and triterpenoids was performed according to Harborne [18]. Total flavonoid content was quantified by the aluminium chloride colorimetric method with quercetin as the reference standard [19]. A quercetin stock solution was prepared by dissolving 10 mg in 10 mL of analytical-grade ethanol and diluted to working standards of 20, 30, 40, 50, 60, 70, and 80 ppm. For each measurement, 1 mL of standard or sample solution was combined with 1 mL of 10% (w/v) aluminium chloride and 1 mL of 10% (w/v) sodium acetate, giving a final reaction volume of 3 mL. The absorption maximum of the quercetin–AlCl₃ complex was established at 419 nm by scanning over 370–450 nm against a reagent blank, and the optimum reaction time was determined to be 15 min from an operating-time profile recorded at 5-min intervals over 30 min. Absorbance was read on a UV-Vis spectrophotometer (Cary 60, Agilent Technologies) in triplicate, yielding the calibration equation y = 0.0072x + 0.3005 (R² = 0.9904). Total flavonoid content was calculated as (c × V × DF)/m, where c is the quercetin-equivalent concentration (mg/L) derived from the calibration curve, V the aliquot volume (L), DF the dilution factor, and m the mass of sample contained in that aliquot (g), and expressed as mg quercetin equivalents per gram of extract (mg QE/g) [20].

### Granule Formulation and Dose

The extract was formulated into granules by wet granulation (Table 1). Polyvinylpyrrolidone was dissolved in 96% ethanol in a mortar, the extract was incorporated and triturated to homogeneity, and the remaining excipients were added and mixed. The mass was passed through a sieve to obtain uniform granules, dried at 50 °C for approximately 15 min to remove residual ethanol, and stored in a tightly closed container. Granules were evaluated for organoleptic properties, flow time, moisture content, compressibility, angle of repose, dissolution time, and pH against established acceptance criteria before use. The formulation is identical to Formula III previously reported by Fadilah et al.[13], which exhibited the strongest xanthine oxidase inhibition among the formulations tested (IC₅₀ 47.18 µg/mL). Because the formulation contains 5% (w/w) extract, 100 mg of granules corresponds to 5 mg of extract; all doses are expressed as granule mass, with extract-equivalent doses given in Table 2. Granules were freshly suspended in 1% sodium carboxymethylcellulose (CMC-Na) immediately before administration and given by oral gavage at 10 mL/kg body weight.

**Table 1.**
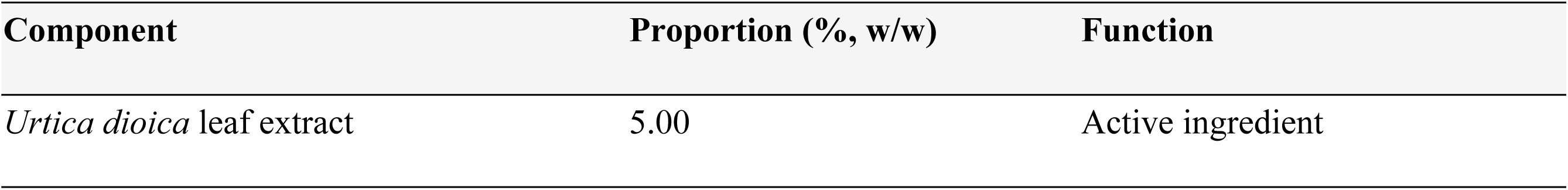
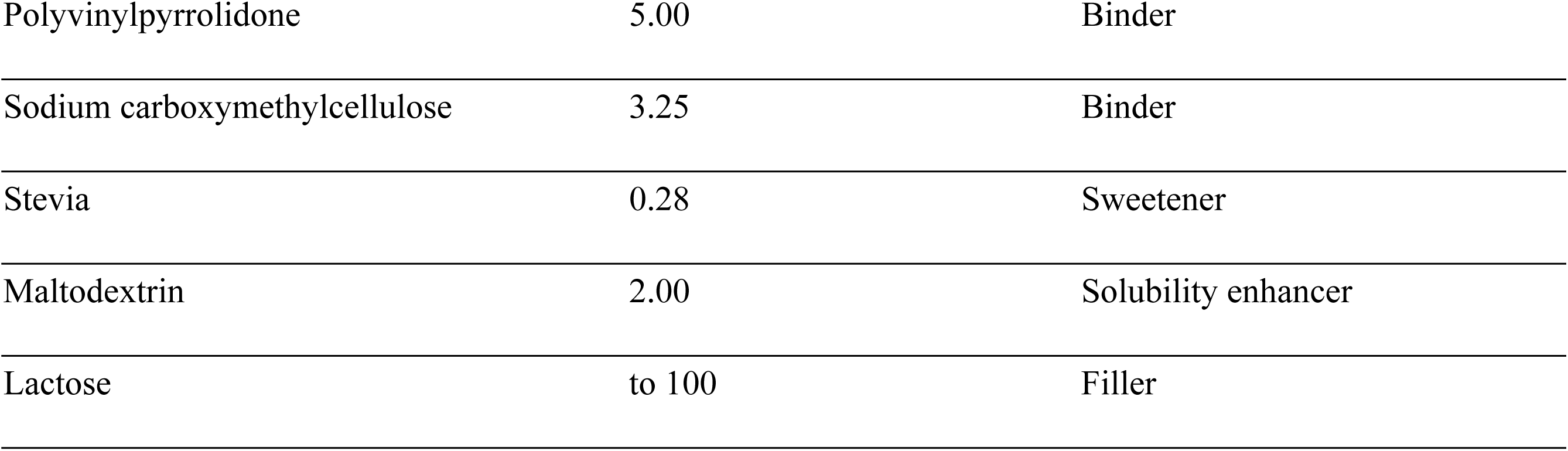
Composition of the *Urtica dioica* leaf extract granule formulation [13].

**Table 2.**
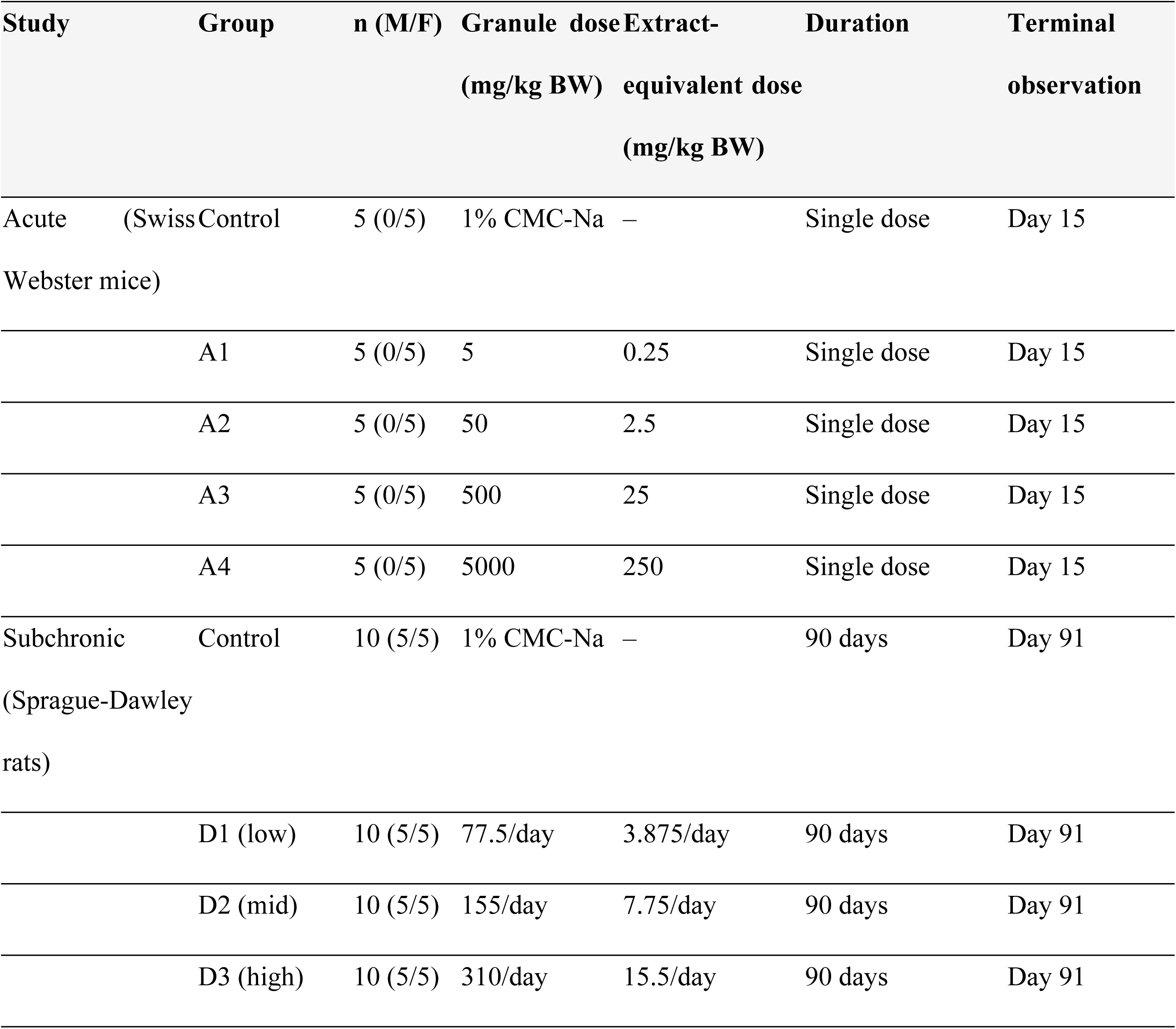
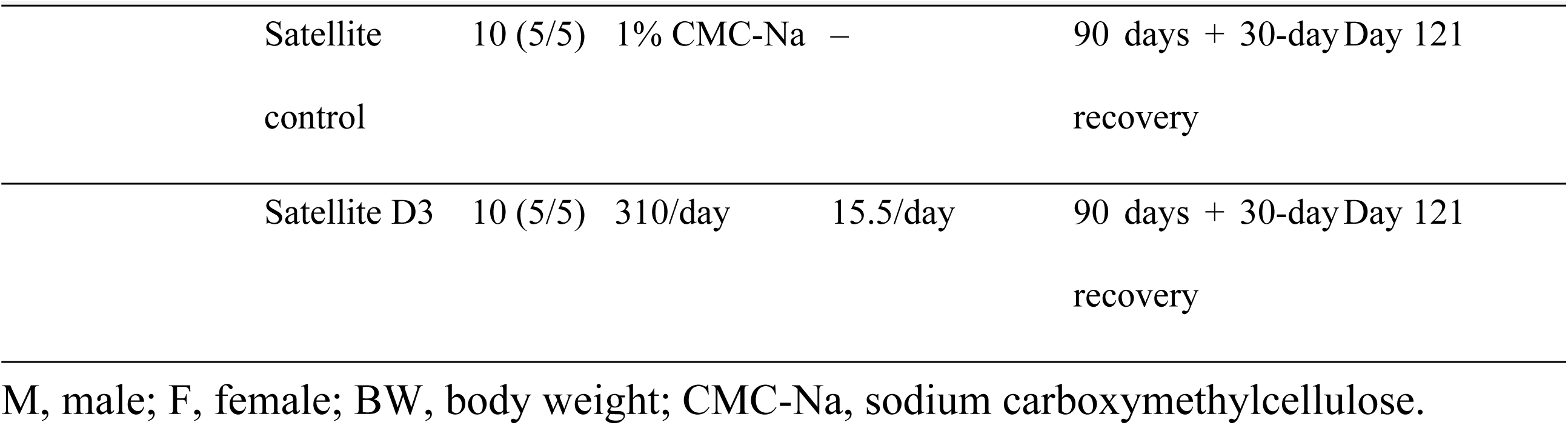
Design of the acute and subchronic oral toxicity studies of *Urtica dioica* leaf extract granules.

### Animal and Husbandry

Female Swiss Webster mice (2–3 months old, 20–35 g) were used for the acute study and Sprague-Dawley rats of both sexes (6–8 weeks old, 150–250 g) for the subchronic study. Animals were supplied by CV. Rumah Tikus Fahlevi (Tasikmalaya, Indonesia) and housed in the Pharmacology Laboratory, Universitas Perjuangan Tasikmalaya, under specific pathogen-free conditions in plastic cages with softwood bedding (five animals per cage) at 23–26 °C and approximately 60% relative humidity under a 12-h light/dark cycle (lights on 08:00–20:00), with purified water and standard pelleted diet available *ad libitum*. Females were nulliparous and non-pregnant. Animals were acclimatised for one week and allocated to the experimental groups. The study was approved by the Health Research Ethics Committee of Universitas Bhakti Tunas Husada Tasikmalaya (No. 057-01/E.01/KEPK-BTH/II/2026, approved 18 May 2026) and conducted in accordance with the Guide for the Care and Use of Laboratory Animals.

### Study Design and Dose Selection

Both studies followed Regulation of the Indonesian Food and Drug Authority (BPOM) No. 10 of 2022 on Guidelines for *In Vivo* Preclinical Toxicity Testing, the national framework corresponding to OECD Test Guidelines 420 and 408 (Table 2). The acute study used the fixed-dose method in female mice only, including the vehicle control group, as the guideline specifies a single sex for the alternative acute methods and identifies females as the more sensitive sex for LD₅₀ estimation. The subchronic study included five males and five females per main and satellite group; the guideline specifies a minimum of ten animals of each sex for the control group, and the use of five per sex represents a deviation adopted to limit animal numbers. Subchronic dose levels were derived from the pharmacologically effective dose of the same granule formulation established in our previous antihyperuricemic study in potassium oxonate-induced mice [21], after interspecies conversion to the rat; the low dose corresponds to the converted effective dose, and the mid and high doses were set at two- and four-fold multiples to permit detection of a dose-response relationship.

### Acute Oral Toxicity Study

After approximately 8 h of fasting with free access to water, mice received a single oral dose. Mortality and general condition were monitored continuously for the first 4 h and daily thereafter for 14 days. Clinical signs were recorded at 2 and 24 h and comprised motor activity, sedation, righting reflex, catalepsy, tremor, salivation, pupil diameter, excretion, piloerection, Straub tail response, Haffner response, pinna reflex, and respiratory changes. Body weight was recorded at baseline, 2 h, 24 h, and on days 7, 10, and 14, and rectal temperature at baseline, 2 h, and 24 h; food and water consumption were not recorded in this phase [9]. Surviving animals were euthanised on day 15 and examined macroscopically. As no mortality occurred at any dose, a numerical LD_₅₀_ could not be estimated by the Thompson-Weil method and is reported as exceeding 5000 mg/kg granules [22].

### Subchronic Study

Rats received vehicle or granules by oral gavage daily for 90 consecutive days. Satellite groups received vehicle or the high dose for the same period and were then maintained untreated for a 30-day recovery period to assess reversibility and delayed toxicity. Animals were observed twice daily for mortality and clinical signs, and body weight was recorded weekly on 12 occasions over the dosing period, with the administered volume recalculated at each weighing. Body weight is reported both as absolute values and as percentage change from the first weighing, the latter serving as the primary growth endpoint because initial body weight was not matched across groups.

### Terminal Procedure and End Point Analyses

After 12 h of fasting, main-group animals were euthanised by inhalation of chloroform vapour on day 91 and satellite animals on day 121 under anaesthesia with ketamine (80 mg/kg) and xylazine (10 mg/kg) intraperitoneally [23]. Blood was collected via the retro-orbital sinus; approximately 0.5 mL was transferred into heparinised tubes and analysed on an automated haematology analyser (Sysmex XP-100, Sysmex Corporation, Kobe, Japan) for erythrocyte count, haemoglobin, haematocrit, erythrocyte indices, platelet count and indices, and total and differential leukocyte counts. Blood for serum biochemistry was collected without anticoagulant, clotted for 60 min at room temperature, and centrifuged at 3,500 rpm (1,370 × g, rotor radius 10 cm) for 15 min before analysis on an automated analyser (BS-860, Mindray, China) for blood urea nitrogen (mg/dL), creatinine (mg/dL), aspartate aminotransferase (U/L), and alanine aminotransferase (U/L).

At necropsy, five animals per group were randomly selected for organ weight determination and histopathological examination; both endpoints were obtained from this same subset. The liver, heart, kidneys, and lungs of the selected animals were excised, rinsed in cold phosphate-buffered saline (PBS), and weighed. Tissues were fixed in 4% buffered formalin (pH 7.4), processed routinely, sectioned at 5 µm, and stained with haematoxylin and eosin. Degeneration, necrosis, inflammation, and haemorrhage were graded on a scale of 0–4 (0, no change; 1, <25%; 2, 25–50%; 3, 50–75%; 4, >75%), with fields treated as subsamples of a single slide rather than independent biological replicates. Slides were coded and scored blind to treatment allocation, and all scores were reviewed and confirmed by a veterinary pathologist [24,25].

#### Statistical Analysis and Criteria for Toxicological Relevance

Data are expressed as mean ± standard deviation and analysed in SPSS version 25. Normality and homogeneity of variance were assessed by the Shapiro-Wilk and Levene tests; homogeneous data were analysed by one-way ANOVA followed by the least significant difference test, and non-homogeneous data by the Kruskal-Wallis test followed by the Mann-Whitney U test. Satellite groups were compared with the satellite control by the Mann-Whitney U test. Significance was set at p < 0.05. Because organ weight and histopathological endpoints were obtained from a sampled subset of five animals per group, these analyses are reported as supportive and exploratory data rather than as confirmatory evidence, and histopathological scores were summarised descriptively without inferential testing. A change reaching statistical significance was considered treatment-related only if it followed a dose-response pattern, fell outside the reference range for the strain, correlated with organ weight or histopathological findings, or persisted after the recovery period [26].

## Results

### Extract and granule characteristics

Remaceration of 500 g of dried leaf powder yielded 16.25 % (w/w) of viscous ethanolic extract. Qualitative screening detected alkaloids, flavonoids, saponins, triterpenoids, tannins, and polyphenols in both the dried powder and the extract. The granule batch administered to the animals contained 5% (w/w) extract and met all pre-specified quality criteria. The granules had a moisture content of 4.8%, a compressibility index of 9.74%, an angle of repose of 30.54°, a dissolution time of 51.41 s, and a pH of 6.94. The total flavonoid content of the extract was 13.38 ± 0.05 mg QE/g of extract.

### Acute Oral Toxicity: Mortality and Clinical Signs

No mortality occurred in any group during the 14-day observation period, including at the limit dose of 5000 mg/kg granules (250 mg/kg extract). As no deaths occurred at the highest dose tested, the LD₅₀ could not be estimated numerically by the Thompson-Weil method and is reported as exceeding 5000 mg/kg body weight.

Transient clinical signs were recorded in both treated and control animals (Table 3). At 2 h, the proportion of animals showing sedation differed significantly among groups (p = 0.001), whereas tremor, piloerection, and respiratory changes did not. At 24 h, between-group differences were also detected for sedation, tremor, and respiratory changes. The incidence of these signs did not increase consistently with dose: sedation occurred in 5/5 animals at 5 mg/kg but in 3/5 at 5000 mg/kg, and tremor was observed in one control animal at 2 h. All signs resolved within the observation period and were not accompanied by mortality.

**Table 3.**
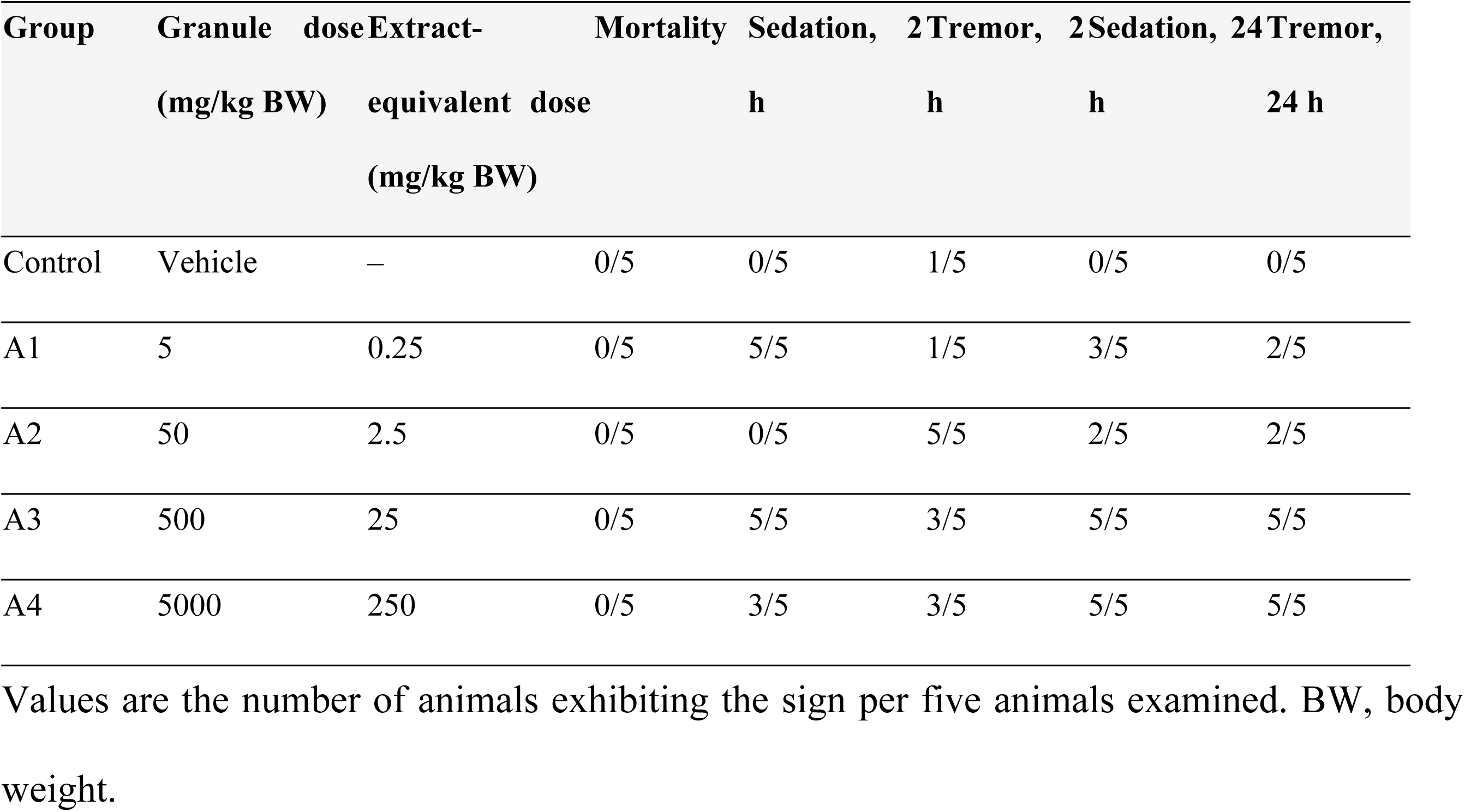
Mortality and incidence of selected clinical signs in female mice after a single oral dose.

| Group | Granule dose<br>(mg/kg BW) | Extract-equivalent dose<br>(mg/kg BW) | Mortality | Sedation, 2 h | Tremor, 2 h | Sedation, 24 h | Tremor, 24 h |
| --- | --- | --- | --- | --- | --- | --- | --- |
| Control | Vehicle | – | 0/5 | 0/5 | 1/5 | 0/5 | 0/5 |
| A1 | 5 | 0.25 | 0/5 | 5/5 | 1/5 | 3/5 | 2/5 |
| A2 | 50 | 2.5 | 0/5 | 0/5 | 5/5 | 2/5 | 2/5 |
| A3 | 500 | 25 | 0/5 | 5/5 | 3/5 | 5/5 | 5/5 |
| A4 | 5000 | 250 | 0/5 | 3/5 | 3/5 | 5/5 | 5/5 |

Values are the number of animals exhibiting the sign per five animals examined. BW, body weight.

### Acute Study: Body Weight and Body Temperature

Body weight is among the most sensitive general indicators of systemic toxicity, and a loss exceeding 10% of the initial value is conventionally regarded as adverse [26]. No such loss occurred. All groups of mice gained weight over the 14-day observation period, from 8.4% in the vehicle control to 14.3% at 5000 mg/kg, with no significant between-group difference at any time point. The transient reduction at 2 h affected treated and control animals alike and had reversed by 24 h, consistent with the overnight fast and gavage procedure rather than a compound-related effect.

Rectal temperature followed the same pattern. Group means did not differ at baseline (p = 0.264) or at 24 h (p = 0.871), and the only significant difference occurred at 2 h in the 500 mg/kg group, whose mean was 0.86 °C above control (36.06 ± 0.15 versus 35.20 ± 0.82 °C). The control group showed the widest dispersion at that time point, one animal recording 33.9 °C. Based on the criteria described in Section 2.10, the relationship of this isolated finding to treatment remained uncertain, and its toxicological significance appeared limited [26].

### 3.4. Subchronic study: body weight

Body weight was recorded on 12 occasions over the 90-day study period and is shown by sex in Table 4. All groups of both sexes gained weight over the study period. Mean percentage weight gain did not differ significantly among groups in either sex (females p = 0.080; males p = 0.087). In pairwise comparison, the female D3 group showed lower percentage weight gain than the female control group (0.7 ± 1.7 versus 5.7 ± 3.3%; p = 0.032) whereas the male D3 group gained more than the male control (7.6 ± 1.6 versus 3.8 ± 1.1%; p = 0.008); the two sexes thus differed in direction, and neither change exceeded the 10% threshold conventionally regarded as adverse. Interpretation of absolute weights is constrained by baseline imbalance, since initial body weight did not differ significantly among groups in either sex (females p = 0.080; males p = 0.080), although numerical differences were present.

**Table 4.** Body weight of rats during the repeated-dose study.

| Sex | Group | n | Initial body weight<br>(g) | Final body weight<br>(g) | Weight gain (%) |
| --- | --- | --- | --- | --- | --- |
| Female | Control | 5 | 175.2 ± 9.5 | 185.2 ± 10.4 | 5.7 ± 3.3 |
|  | D1 | 5 | 182.6 ± 4.0 | 195.2 ± 7.4 | 6.9 ± 2.3 |
|  | D2 | 5 | 184.4 ± 24.6 | 194.0 ± 20.9 | 5.5 ± 3.9 |
|  | D3 | 5 | 197.2 ± 10.4 | 198.4 ± 8.1 | 0.7 ± 1.7* |
|  | Satellite control | 5 | 188.2 ± 4.9 | 190.2 ± 5.9 | 1.1 ± 3.0 |
|  | Satellite D3 | 5 | 185.4 ± 4.2 | 193.6 ± 6.8 | 4.4 ± 2.4 |
|  | Kruskal-Wallis p |  | 0.080 | 0.287 | 0.080 |
| Male | Control | 5 | 179.4 ± 5.8 | 186.0 ± 5.4 | 3.8 ± 1.1 |
|  | D1 | 5 | 199.4 ± 15.6 | 209.2 ± 13.1 | 5.0 ± 1.6 |
|  | D2 | 5 | 203.4 ± 17.1 | 213.8 ± 17.7 | 5.1 ± 2.5 |
|  | D3 | 5 | 192.0 ± 7.6 | 205.0 ± 6.0 | 7.6 ± 1.6* |
|  | Satellite control | 5 | 196.8 ± 7.0 | 204.8 ± 7.8 | 4.1 ± 0.9 |
|  | Satellite D3 | 5 | 189.6 ± 10.1 | 195.4 ± 6.7 | 3.1 ± 2.3 |
|  | Kruskal-Wallis p |  | 0.080 | 0.009 | 0.087 |

Values are mean ± SD. \* $p < 0.05$ versus the corresponding control group (Mann-Whitney U test).

### Subchronic Study: Organ Weights

Absolute organ weights are presented separately by sex in Tables 5 and 6. In females, liver weight differed significantly among groups (p = 0.031); mean liver weight was higher in D1 than in the control group (8.52 ± 0.58 versus 6.60 ± 1.54 g; p = 0.032), and the satellite D3 group was higher than the satellite control (8.63 ± 0.30 versus 7.49 ± 0.28 g; p = 0.012). The difference did not follow a dose-response pattern, since liver weight at the mid and high doses did not differ from control (p = 0.095 for both). Lung weight also differed among female groups (p = 0.019), but no treated group differed from the control in pairwise comparison (p = 0.173– 0.421), the overall difference arising between the D2 and D3 groups. Heart and kidney weights did not differ (p = 0.735 and 0.137). In males, no organ weight differed significantly among groups (p = 0.067–0.922).

**Table 5.**
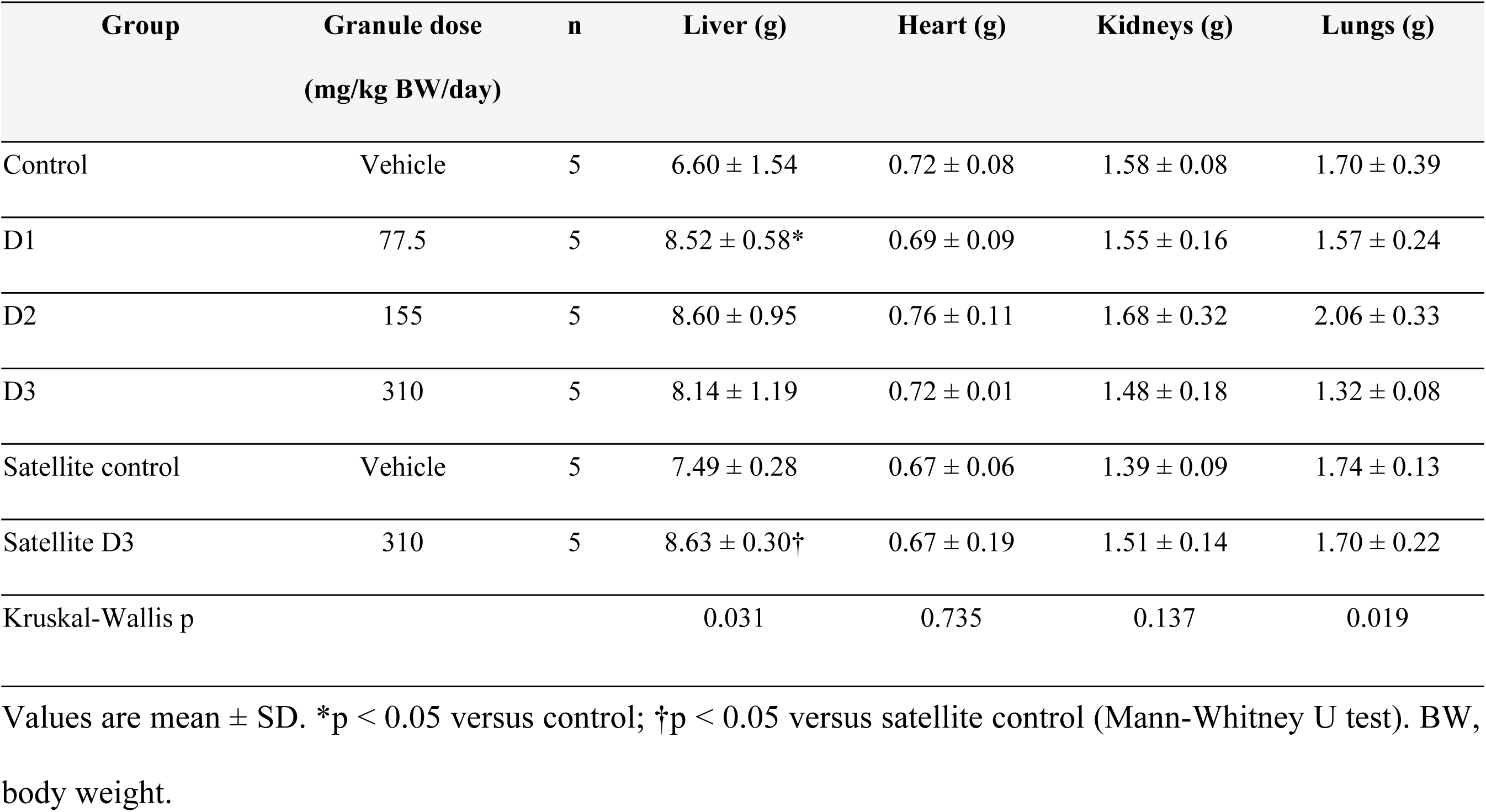
Absolute organ weights of female rats at the end of dosing and after the recovery period.

**Table 6.** Absolute organ weights of male rats at the end of dosing and after the recovery period.

| Group | Granule dose<br>(mg/kg BW/day) | n | Liver (g) | Heart (g) | Kidneys (g) | Lungs (g) |
| --- | --- | --- | --- | --- | --- | --- |
| Control | Vehicle | 5 | $9.15 \pm 1.06$ | $0.86 \pm 0.12$ | $1.80 \pm 0.23$ | $1.38 \pm 0.39$ |
| D1 | 77.5 | 5 | $9.66 \pm 0.63$ | $0.87 \pm 0.11$ | $1.95 \pm 0.17$ | $1.58 \pm 0.07$ |
| D2 | 155 | 5 | $9.49 \pm 0.43$ | $0.94 \pm 0.05$ | $1.95 \pm 0.11$ | $1.89 \pm 0.41$ |
| D3 | 310 | 5 | $9.78 \pm 0.77$ | $0.92 \pm 0.09$ | $1.96 \pm 0.18$ | $1.65 \pm 0.15$ |
| Satellite control | Vehicle | 5 | 9.45 ± 0.83 | 1.01 ± 0.09 | 2.36 ± 0.38 | 1.50 ± 0.07 |
| Satellite D3 | 310 | 5 | 9.60 ± 0.80 | 0.94 ± 0.21 | 2.03 ± 0.17 | 1.71 ± 0.18 |
| Kruskal-Wallis p |  |  | 0.922 | 0.465 | 0.067 | 0.083 |

Values are mean ± SD. BW, body weight.

### Haematology and Serum Biochemistry

Haematological and biochemical values are summarised in Table 7. In females, leukocyte count was significantly higher in D3 than in the control (8.3 versus 2.8 × 10³/µL; p < 0.05) and was lower in the D3 satellite group (5.7 × 10³/µL) than in the satellite control (7.0 × 10³/µL). Alanine aminotransferase was significantly lower in D2 than in the control (63 versus 71 U/L; p < 0.05), a change opposite in direction to that expected for hepatocellular injury. Erythrocyte parameters varied without a dose-related pattern, with haemoglobin, erythrocyte count, and haematocrit lowest in D2. Platelet count in the D3 satellite group (220 × 10³/µL) was lower than in the satellite control (239 × 10³/µL). In males, no parameter differed significantly from control. Aspartate aminotransferase was above the reference range for the strain in every group of both sexes, including the vehicle controls (females 137-287 U/L; males 209–225U/L).

**Table 7.** Haematological and serum biochemical parameters of rats.

| Parameter | Unit | Control | D1 | D2 | D3 | Satellite control | Satellite D3 |
| --- | --- | --- | --- | --- | --- | --- | --- |
| Female |  |  |  |  |  |  |  |
| Haemoglobin | g/dL | 15.1 ± 1.71 | 13.0 ± 1.19 | 9.3 ± 1.65 | 13.5 ± 1.00 | 13.1 ± 0.43 | 14.6 ± 1.32 |
| Leukocytes | $\times 10^3/\mu\text{L}$ | 2.8 ± 2.66 | 2.8 ± 0.12 | 3.7 ± 2.56 | 8.3 ± 4.33* | 7.0 ± 2.88 | 5.7 ± 0.11 |
| Erythrocytes | $\times 10^6/\mu\text{L}$ | 5.0 ± 0.25 | 6.4 ± 0.44 | 4.1 ± 0.32 | 6.5 ± 0.89 | 5.9 ± 0.21 | 6.5 ± 0.32 |
| Haematocrit | % | 29 ± 0.12 | 37 ± 4.21 | 25 ± 2.34 | 35 ± 1.10 | 34 ± 1.98 | 37 ± 3.21 |
| Platelets | $\times 10^3/\mu\text{L}$ | 260 ± 2.22 | 188 ± 2.10 | 387 ± 5.10 | 312 ± 3.65 | 239 ± 6.43 | 220 ± 3.10 |
| MCV | fL | 58 ± 8.65 | 57 ± 1.04 | 59 ± 2.90 | 54 ± 0.41 | 58 ± 2.11 | 57.6 ± 3.13 |
| MCH | pg | 30.2 ± 1.14 | 28.9 ± 0.21 | 30.4 ± 1.24 | 30.1 ± 2.55 | 32 ± 1.05 | 30.2 ± 0.21 |
| MCHC | g/dL | 52.1 ± 3.21 | 54 ± 2.65 | 53 ± 6.21 | 53.2 ± 1.22 | 50 ± 6.00 | 50 ± 0.11 |
| Blood urea nitrogen | mg/dL | 61 ± 1.09 | 52 ± 1.07 | 55 ± 4.09 | 61 ± 2.12 | 47 ± 2.11 | 47 ± 1.98 |
| Creatinine | mg/dL | 0.5 ± 0.11 | 0.4 ± 0.11 | 0.6 ± 0.34 | 0.6 ± 0.23 | 0.6 ± 0.09 | 0.5 ± 0.12 |
| AST | U/L | 201 ± 2.03 | 187 ± 3.76 | 262 ± 3.67 | 287 ± 3.43 | 243 ± 3.30 | 137 ± 4.31 |
| ALT | U/L | 71 ± 1.07 | 66 ± 0.11 | 63 ± 9.22* | 77 ± 9.50 | 90 ± 5.88 | 73 ± 0.11 |
| Male |  |  |  |  |  |  |  |
| Haemoglobin | g/dL | 13.5 ± 0.8 | 14.8 ± 0.7 | 11.3 ± 0.9 | 14.6 ± 1.0 | 12.0 ± 0.6 | 12.7 ± 0.8 |
| Leukocytes | ×10<sup>3</sup>/μL | 7.2 ± 1.4 | 5.6 ± 1.6 | 5.9 ± 1.2 | 6.1 ± 1.8 | 5.4 ± 1.3 | 4.8 ± 1.5 |
| Erythrocytes | ×10<sup>6</sup>/μL | 8.0 ± 0.5 | 7.4 ± 0.4 | 7.9 ± 0.6 | 7.1 ± 0.5 | 8.3 ± 0.4 | 9.0 ± 0.6 |
| Haematocrit | % | 43.5 ± 2.8 | 44.3 ± 2.5 | 42.9 ± 3.1 | 43.8 ± 3.0 | 45.0 ± 2.4 | 44.1 ± 2.9 |
| Platelets | ×10<sup>3</sup>/μL | 780 ± 10.5 | 815 ± 9.2 | 760 ± 11.8 | 830 ± 12.5 | 795 ± 8.8 | 805 ± 11.0 |
| MCV | fL | 58.4 ± 2.1 | 58.0 ± 1.8 | 59.3 ± 2.4 | 60.1 ± 2.0 | 54.2 ± 1.7 | 58.4 ± 2.2 |
| MCH | pg | 30.1 ± 0.8 | 31.8 ± 0.7 | 30.1 ± 0.9 | 30.2 ± 0.8 | 30.1 ± 0.6 | 32.1 ± 0.9 |
| MCHC | g/dL | 48.1 ± 1.1 | 53.4 ± 1.0 | 51.3 ± 1.3 | 49.3 ± 1.2 | 48.3 ± 0.9 | 53.3 ± 1.1 |
| Blood urea nitrogen | mg/dL | 60.5 ± 3.2 | 69.2 ± 2.8 | 58.0 ± 3.5 | 60.1 ± 3.8 | 59.0 ± 2.6 | 59.6 ± 3.1 |
| Creatinine | mg/dL | 0.48 ± 0.08 | 0.50 ± 0.07 | 0.47 ± 0.09 | 0.53 ± 0.10 | 0.49 ± 0.06 | 0.51 ± 0.08 |
| AST | U/L | 212 ± 9.0 | 218 ± 2.5 | 209 ± 1.9 | 225 ± 2.8 | 215 ± 2.0 | 220 ± 2.4 |
| ALT | U/L | 68 ± 2.76 | 54 ± 3.31 | 57 ± 7.0 | 64 ± 4.12 | 49 ± 8.0 | 43 ± 9.0 |

Values are mean ± SD. \*p < 0.05 versus the corresponding control group. AST, aspartate aminotransferase; ALT, alanine aminotransferase; MCV, mean corpuscular volume; MCH, mean corpuscular haemoglobin; MCHC, mean corpuscular haemoglobin concentration.

### Histopathology

Microscopic findings are summarised in Table 8 and illustrated in Figs 3 and 4. In the liver, mean scores for degeneration, necrosis, and inflammation ranged from 0.2 to 1.2, the highest being necrosis in D3 (1.2), corresponding to mild change on the scale used. Renal scores remained between 0.0 and 0.8 with no consistent increase after the recovery period. In the heart, the D3 satellite group recorded a degeneration score of 1.2, higher than both the main D3 group (0.4) and the satellite control (0.2). In the lung, inflammation and haemorrhage scores in the D3 satellite group were 1.8 each, compared with 1.0 and 0.4 in the satellite control. Because scoring was based on five microscopic fields from a single slide per group, these differences are presented descriptively and were not tested inferentially.

**Fig 1.**
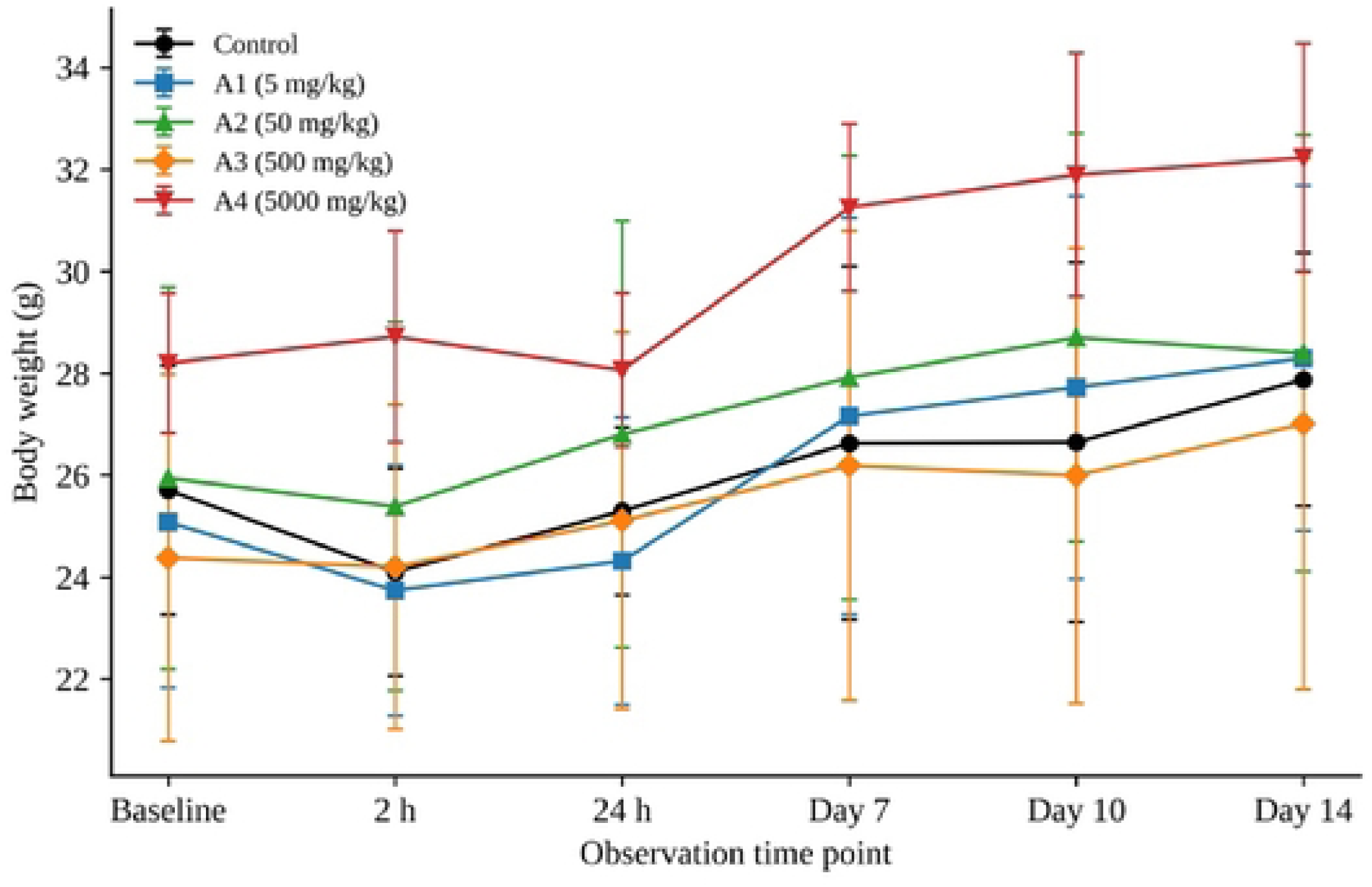
Body weight of female mice over the 14-day observation period following a single oral dose of *Urtica dioica* leaf extract granules. Points represent group means and error bars represent standard deviations.

**Fig 2.**
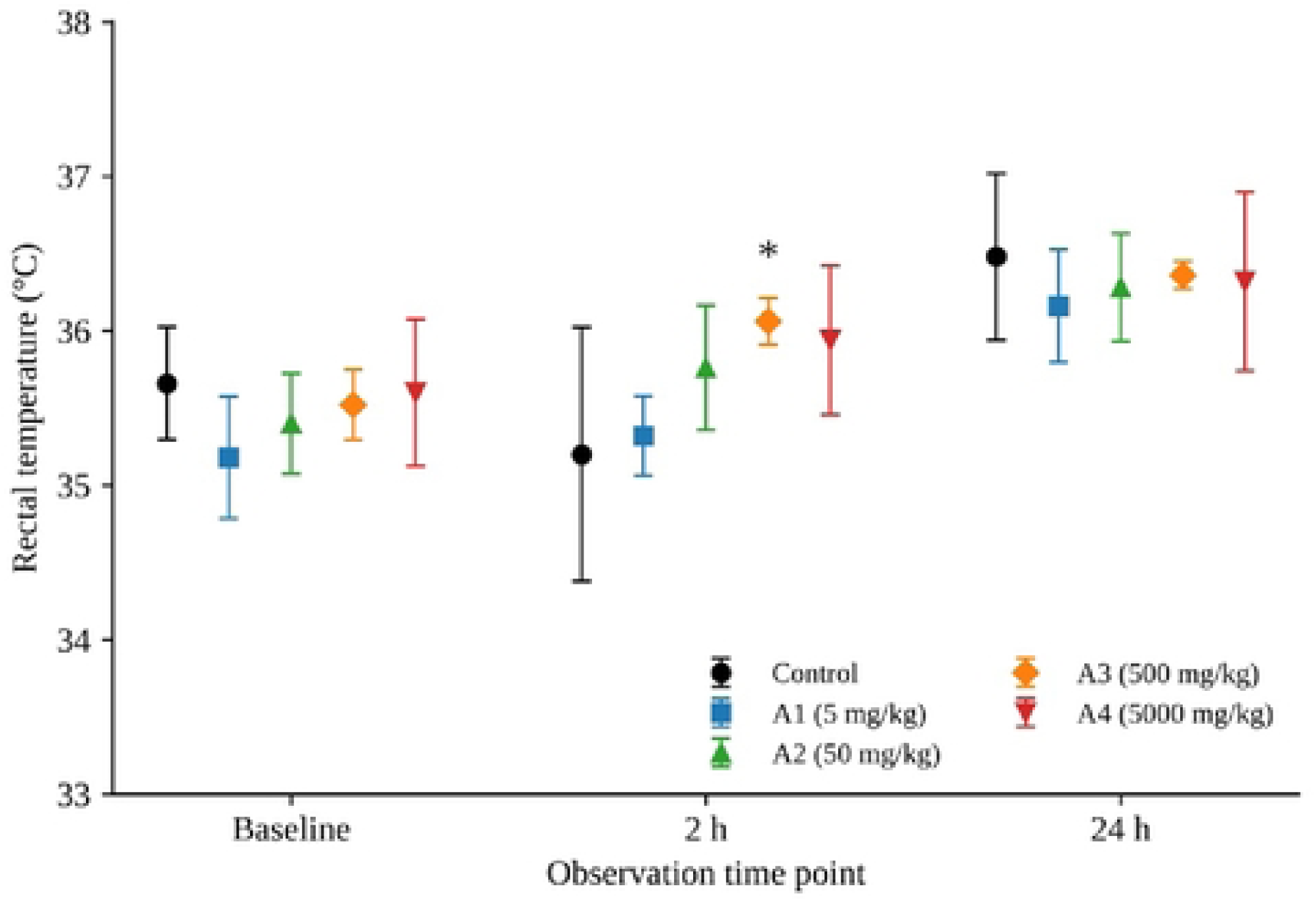
Body temperature of female mice over the 14-day observation period following a single oral dose of *Urtica dioica* leaf extract granules. Points represent group means and error bars represent standard deviations.

**Fig 3.**
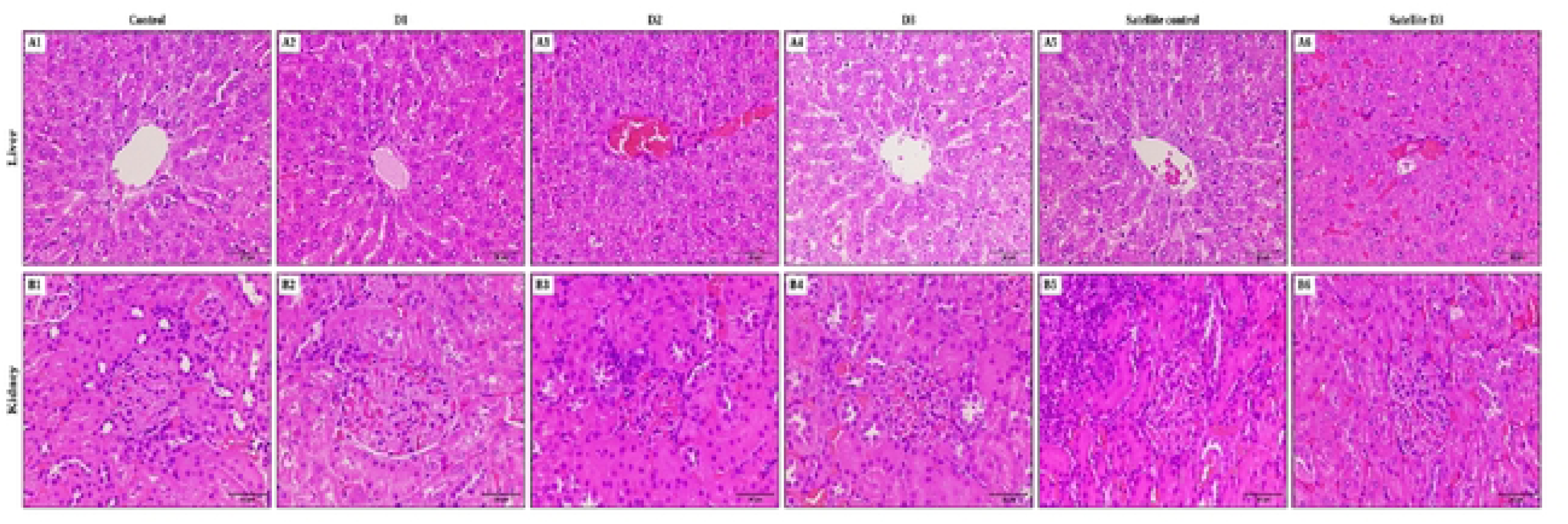
Representative photomicrographs of liver (A1–A6) and kidney (B1–B6) from rats treated with *Urtica dioica* leaf extract granules for 90 days and from the recovery groups. Columns correspond to the control, D1 (77.5 mg/kg/day), D2 (155 mg/kg/day), D3 (310 mg/kg/day), satellite control, and satellite D3 groups. Haematoxylin and eosin, ×400; scale bar = 50 µm.

**Fig 4.**
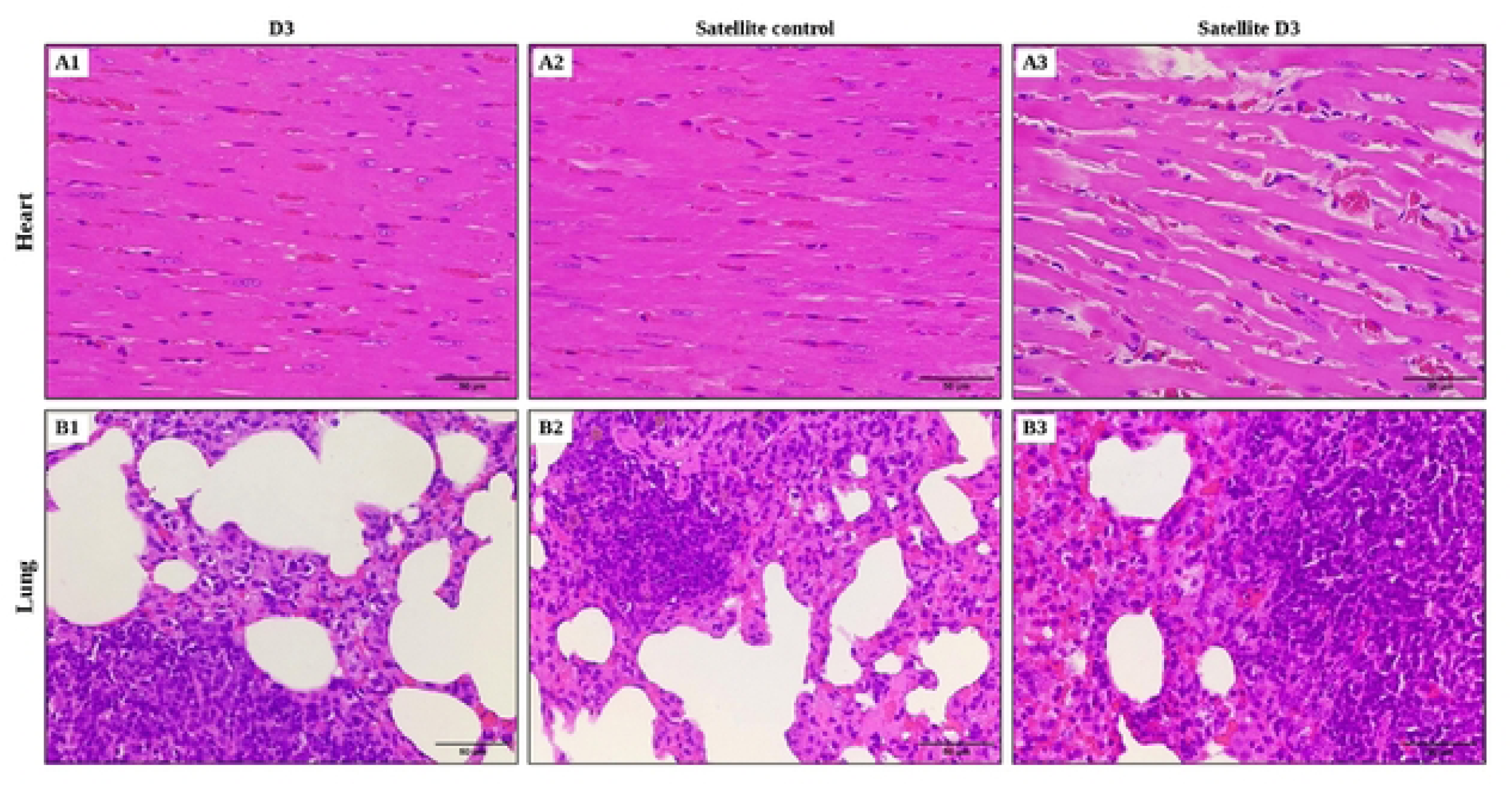
Representative photomicrographs of heart (A1–A3) and lung (B1–B3) from the D3 (310 mg/kg/day), satellite control, and satellite D3 groups. Haematoxylin and eosin, ×400; scale bar = 50 µm.

**Table 8.**
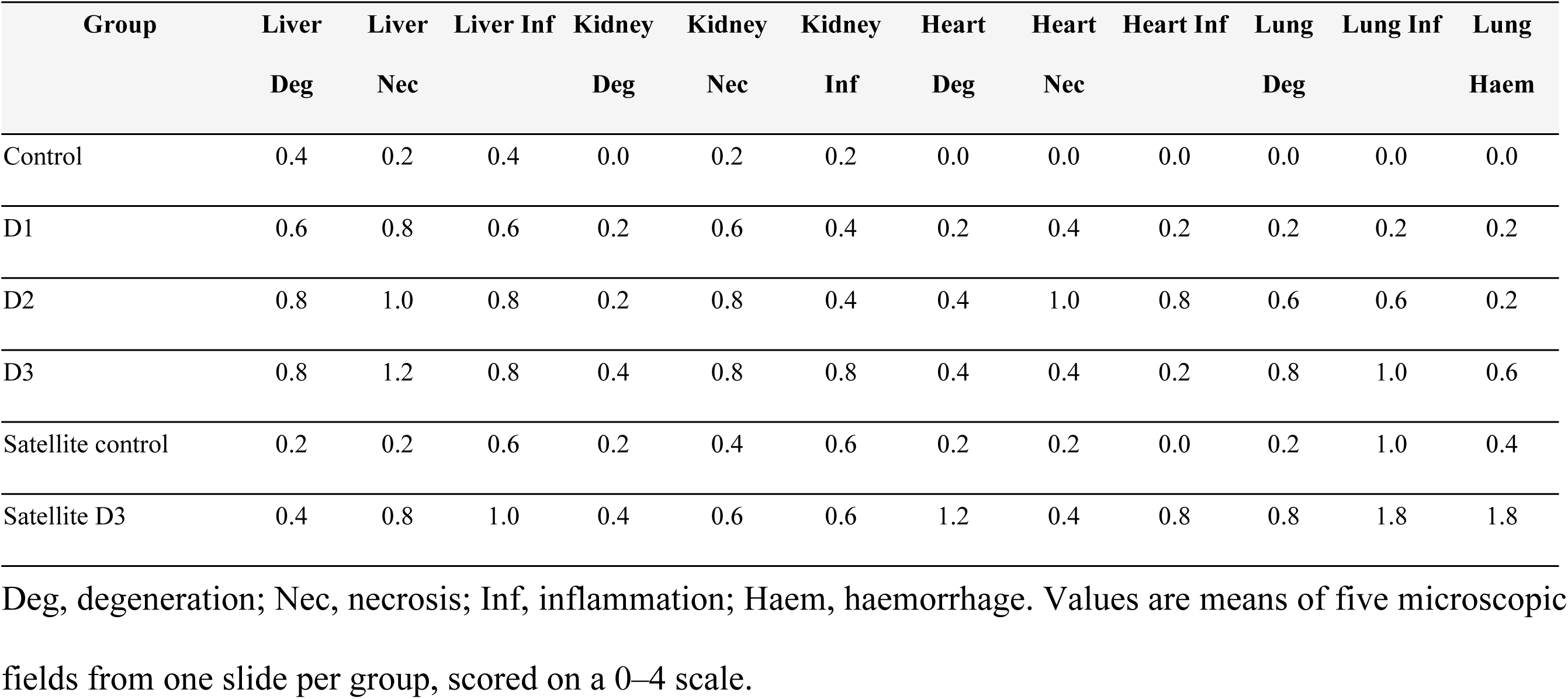
Mean histopathological scores at the end of dosing and after the recovery period.

| Group | Liver<br>Deg | Liver<br>Nec | Liver Inf | Kidney<br>Deg | Kidney<br>Nec | Kidney<br>Inf | Heart<br>Deg | Heart<br>Nec | Heart Inf | Lung<br>Deg | Lung Inf | Lung<br>Haem |
| --- | --- | --- | --- | --- | --- | --- | --- | --- | --- | --- | --- | --- |
| Control | 0.4 | 0.2 | 0.4 | 0.0 | 0.2 | 0.2 | 0.0 | 0.0 | 0.0 | 0.0 | 0.0 | 0.0 |
| D1 | 0.6 | 0.8 | 0.6 | 0.2 | 0.6 | 0.4 | 0.2 | 0.4 | 0.2 | 0.2 | 0.2 | 0.2 |
| D2 | 0.8 | 1.0 | 0.8 | 0.2 | 0.8 | 0.4 | 0.4 | 1.0 | 0.8 | 0.6 | 0.6 | 0.2 |
| D3 | 0.8 | 1.2 | 0.8 | 0.4 | 0.8 | 0.8 | 0.4 | 0.4 | 0.2 | 0.8 | 1.0 | 0.6 |
| Satellite control | 0.2 | 0.2 | 0.6 | 0.2 | 0.4 | 0.6 | 0.2 | 0.2 | 0.0 | 0.2 | 1.0 | 0.4 |
| Satellite D3 | 0.4 | 0.8 | 1.0 | 0.4 | 0.6 | 0.6 | 1.2 | 0.4 | 0.8 | 0.8 | 1.8 | 1.8 |

Deg, degeneration; Nec, necrosis; Inf, inflammation; Haem, haemorrhage. Values are means of five microscopic fields from one slide per group, scored on a 0–4 scale.

## Discussion

Oral administration of *Urtica dioica* leaf extract granules produced no mortality at a single dose of 5000 mg/kg and no consistent, dose-related adverse change after 13 weeks (90 days) of daily dosing in rats of both sexes. The isolated findings described below, together with control values outside the expected ranges for the strain and an imbalance in body weight at allocation, mean that a definitive no-observed-adverse-effect level cannot yet be established from this dataset.

The absence of mortality at the limit dose places the granules in the practically non-toxic range under the Globally Harmonized System, and the LD₅₀ is reported as exceeding 5000 mg/kg granules, equivalent to 250 mg/kg extract. All groups of mice gained weight over the 14 days observation period, and no group approached the 10% loss conventionally regarded as adverse. The transient weight reduction at 2 h affected treated and control animals alike and had reversed by 24 h, consistent with the overnight fast and the gavage procedure rather than with a compound-related effect.

Rectal temperature did not differ among groups at baseline or at 24 h. The only statistically significant difference was observed at 2 h in the 500 mg/kg group; it was absent at the highest dose and was not detected at 24 h. These observations suggest limited toxicological significance, the differing directions of change between sexes complicate interpretation and do not exclude a sex-specific treatment effect.. The greater variability in the control group, including one reading of 33.9 °C, warrants consideration when interpreting this isolated comparison.

In the repeated-dose study, the only significant growth findings occurred at the highest dose and ran in opposite directions in the two sexes: female rats gained less weight than controls while male animals gained more. A treatment-related effect on growth would be expected to act in the same direction in both sexes, and neither change approached the 10% threshold. Interpretation is further limited by baseline imbalance, since initial body weight did not differ significantly among groups in both sexes and the control group of each sex was the lightest at the first weighing. Percentage change was therefore used as the growth endpoint, and the significant difference in final male body weight is attributed to that imbalance rather than to treatment.

Haematological findings followed a similar pattern. Leukocyte count was significantly higher in the female high-dose group than in its control, but the increase was absent at lower doses, was not reproduced in males, and this increase was not observed in the high-dose satellite group at the end of the recovery period. Erythrocyte indices remained stable across all groups and no male parameter differed significantly from control. These findings suggest limited evidence of an adverse haematological effect, although a reversible, female-specific treatment-related increase in leukocyte count cannot be excluded. Isolated statistically significant haematological changes without a consistent dose-related pattern have also been reported in subchronic studies of botanical preparations [27].

The reduction in alanine aminotransferase activity in the female mid-dose group did not represent the enzyme elevation typically associated with hepatocellular injury. Aspartate aminotransferase activity exceeded the cited reference range in all groups of both sexes, including vehicle controls, without a corresponding increase in alanine aminotransferase. This distribution suggests a common background or procedural influence rather than a clear treatment-specific effect. Retro-orbital sampling may contribute to elevated aspartate aminotransferase activity through local tissue injury [28][29,30]; however, the contribution of haemolysis could not be established without documented assessment of sample quality. Creatinine and urea showed no detected treatment-related changes, providing no biochemical evidence of overt renal dysfunction, although subtle renal injury cannot be excluded.

Absolute organ weights showed one finding that warrants comment. In females, liver weight differed significantly among groups, with the low-dose and satellite high-dose groups exceeding their respective controls. The difference did not follow a dose-response pattern, since the mid- and high-dose groups did not differ from control, and the female control group had the lowest mean liver weight of any group. No corresponding difference was found in males. The absence of a consistent dose-related pattern limits attribution of the liver weight differences to treatment.

Histopathological scores remained in the mild range throughout. The findings of greatest interest arose in the high-dose satellite group, where cardiac degeneration and pulmonary inflammation and haemorrhage were higher than in both the corresponding dosing group and the satellite control. These lesions were observed at the recovery necropsy, but their time of onset could not be established because different animals were examined at the end of dosing and recovery. Their relationship to treatment remains uncertain. Pulmonary inflammation in the satellite control suggests a possible background contribution, although a colony-wide respiratory condition was not established. Several limitations qualify these conclusions. Allocation was not stratified by body weight, and initial body weight did not differ significantly among groups in both sexes [30]. The control group size was below that specified by the guideline. Aspartate aminotransferase values lay outside the reference range in all groups including controls, weakening every comparison anchored to those groups. The biochemical panel was limited to four analytes, so alkaline phosphatase, total protein, albumin, bilirubin, glucose, and the lipid profile were unavailable to support the interpretation of hepatic findings. Histopathology was scored from a single slide per group rather than per animal. Finally, the extract was characterised by total flavonoid content alone, without identification of marker compounds, so extrapolation to other extract batches requires caution.

## Conclusion

Under the conditions of this study, a single oral dose of *Urtica dioica* leaf extract granules produced no mortality in female mice up to 5000 mg/kg body weight (250 mg/kg extract-equivalent), giving an LD₅₀ above that limit and placing the formulation in the practically non-toxic range. Clinical signs, body weight, and rectal temperature showed no dose-related change over the 14-day observation period. After 13 weeks (90 days) of repeated oral administration at 77.5, 155, and 310 mg/kg/day, erythrocyte indices, blood urea nitrogen, and creatinine were unchanged in both sexes, no male parameter differed from control, and the isolated changes observed in females, namely leukocytosis at the highest dose and a liver weight difference without dose-response, did not meet the criteria for treatment-related adverse effects.

These findings are consistent with low systemic toxicity but do not yet support a defined no-observed-adverse-effect level. Initial body weight did not differ significantly among groups in both sexes, aspartate aminotransferase values lay outside the strain reference range in all groups including controls, and the cardiac and pulmonary findings recorded only in the high-dose satellite group require confirmation rather than dismissal. A repeat study with weight-stratified allocation, animal-level histopathology, an expanded biochemical panel, and a blood sampling procedure that avoids haemolysis is required before 310 mg/kg/day can be proposed as a NOAEL and translated into a human equivalent dose.

## Supporting Information

S1 File. The dataset encompasses hematological and biochemical parameters, as well as body weight measurements of the mice (XLXS)

## Acknowledgment

The authors thank Universitas Perjuangan Tasikmalaya for institutional support and research facilities.

## References

1. Im PK, Kartsonaki C, Kakkoura MG, Mohamed-Ahmed O, Yang L, Chen Y, et al. Hyperuricemia, gout and the associated comorbidities in China: findings from a prospective study of 0.5 million adults. Lancet Reg Health West Pac. 2025;58. doi:10.1016/j.lanwpc.2025.101572

2. Farquhar H, Gaffo A, Stamp LK. Safety and efficacy of gout treatments in people with renal impairment. Exploration of Musculoskeletal Diseases. 2024;2(5). doi:10.37349/emd.2024.00062

3. Bove M, Cicero AFG, Veronesi M, Borghi C. An evidence-based review on urate-lowering treatments: Implications for optimal treatment of chronic hyperuricemia. Vascular Health and Risk Management. 2017. doi:10.2147/VHRM.S115080

4. Forster A. [Therapy of gout in 2024]. Ther Umsch. 2024;81(5).

5. Li XX, Han M, Wang YY, Liu JP. Chinese herbal medicine for gout: A systematic review of randomized clinical trials. In: Clinical Rheumatology. 2013. doi:10.1007/s10067-013-2274-7

6. Fadilah NN, Susanti S. Aktivitas Antihiperurisemia Ekstrak Tanaman Jelatang (Urtica dioca L.) pada Mencit. Health Information : Jurnal Penelitian. 2020;12(1). doi:10.36990/hijp.vi.193

7. Omar Ghariba A, Mohammed LY. PHYSIOCHEMICAL STANDARDIZATION AND PHYTOCHEMICAL SCREENING OF Urtica dioica L. LEAVES GROWING IN ZAKHO, KURDISTAN REGION, IRAQ. Science Journal of University of Zakho. 2023;11(3). doi:10.25271/sjuoz.2023.11.3.1069

8. Orhan IE, Deniz FSS. Natural Products and Extracts as Xantine Oxidase Inhibitors - A Hope for Gout Disease? Curr Pharm Des. 2020;27(2). doi:10.2174/1381612826666200728144605

9. Bpom RI. Peraturan BPOM No 10 Tahun 2022 Pedoman Uji Toksisitas Praklinik Secara In Vivo. Bpom Ri. 2022.

10. Dar SA, Ganai FA, Yousuf AR, Balkhi MUH, Bhat TM, Sharma P. Pharmacological and toxicological evaluation of Urtica dioica. Pharm Biol. 2013. doi:10.3109/13880209.2012.715172

11. Wube B, Asres K, Gemechu W, Degu S, Girma Y, Seyoum G. Acute and subacute toxicity of ethanol leaf extract of Urtica simensis Hochst. ex A. Rich in rats. Drug Chem Toxicol. 2026;49(2). doi:10.1080/01480545.2025.2527176

12. Li D, Lyu Y, Zhao J, Ji X, Zhang Y, Zuo Z. Accumulation of the Major Components from Polygoni Multiflori Radix in Liver and Kidney after Its Long-Term Oral Administrations in Rats. Planta Med. 2022;88(11). doi:10.1055/a-1585-5991

13. Fadilah NN, Endah SRN, Sumiwi SA. Evaluation of In Vitro Xanthine Oxidase Inhibitory and In Vivo Antihyperuricemia Activities of Urtica dioica L. Extract Granules. Tropical Journal of Natural Product Research. 2026 May 1;10(4):8367–72. doi:10.26538/tjnpr/v10i4.14

14. Food and Drug Administration. Botanical Drug Development - Guidance for industry. U.S. Department of Health and Human Services. 2016.

15. OECD. Test Guideline No. 407 Repeated Dose 28-day Oral Toxicity Study in Rodents. In: OECD guideline for testing of chemicals. 2025.

16. M3(R2); ICH E. ICH M3(R2) - Guidance on non-clinical safety studies for the conduct of human clinical trials and marketing authorization for pharmaceuticals. International Conference on Harmonisation. 2009;3(June).

17. Kemenkes RI. Farmakope Herbal Indonesia Edisi II 2017 Kementrian Kesehatan Republik Iindonesia. Pills and the Public Purse. 2017.

18. Paterson. Phytochemical Methods. A Guide to Modern Techniques of Plant Analysis. Plant Pathol. 1999;48(1). doi:10.1046/j.1365-3059.1999.00318.x

19. Chia-Chi Chang, Ming-Hua Yang, Hwei-Mei Wen, Jiing-Chuan Chern. Estimation of Total Flavonoid Content in Propolis by Two Complementary Colorimetric Methods. J Food Drug Anal. 2002;10(3).

20. Hoswari CN, Br Karo RM, Yudha M. Penentuan kadar total fenolik, total flavonoid, dan uji aktivitas antibakteri ekstrak etanol daun kerai payung (Filicium decipiens) terhadap bakteri Propionibacterium acnes dan Staphylococcus epidermis. Jurnal Prima Medika Sains. 2023;5(1). doi:10.34012/jpms.v5i1.3699

21. Alghamdi YS, Soliman MM, Nassan MA. Impact of Lesinurad and allopurinol on experimental Hyperuricemia in mice: Biochemical, molecular and Immunohistochemical study. BMC Pharmacol Toxicol. 2020;21(1). doi:10.1186/s40360-020-0386-7

22. Saganuwan SA. The new algorithm for calculation of median lethal dose (LD50) and effective dose fifty (ED50) of Micrarus fulvius venom and anti-venom in mice. Int J Vet Sci Med. 2016;4(1). doi:10.1016/j.ijvsm.2016.09.001

23. Giroux MC, Santamaria R, Hélie P, Burns P, Beaudry F, Vachon P. Physiological, pharmacokinetic and liver metabolism comparisons between 3-, 6-, 12- and 18-month-old male sprague dawley rats under ketamine-xylazine anesthesia. Exp Anim. 2016;65(1). doi:10.1538/expanim.15-0039

24. Schafer KA, Eighmy J, Fikes JD, Halpern WG, Hukkanen RR, Long GG, et al. Use of Severity Grades to Characterize Histopathologic Changes. Toxicol Pathol. 2018;46(3). doi:10.1177/0192623318761348

25. Junsai T, Tangpanithandee S, Srimangkornkaew P, Suknuntha K, Vivithanaporn P, Khemawoot P. Sub-chronic oral toxicity of a water-soluble extract of Centella asiatica (Centell-S) in Wistar rats. Food and Chemical Toxicology. 2024;185. doi:10.1016/j.fct.2024.114509

26. Lewis RW, Billington R, Debryune E, Gamer A, Lang B, Carpanini F. Recognition of Adverse and Nonadverse Effects in Toxicity Studies. Toxicol Pathol. 2002;30(1). doi:10.1080/01926230252824725

27. Madireddy RK, Alluri KV, Somepalli V, Golakoti T, Sengupta K. Toxicological Assessments of a Proprietary Blend of Punica granatum Fruit Rind and Theobroma cacao Seed Extracts: Acute, Subchronic, and Genetic Toxicity Studies. J Toxicol. 2022;2022. doi:10.1155/2022/3903943

28. Yeganeh BS, Soleimani N, Zare S, Mohammadzadeh S, Amiri F, Esmaeili MJ, et al. The Effect of Hemolysis on Biochemical Tests: Determining Interference Cutoffs and Managing Hemolytic Samples. J Anal Methods Chem. 2026;2026(1). doi:10.1155/jamc/9307894

29. Sharma A, Fish BL, Moulder JE, Medhora M, Baker JE, Mader M, et al. Safety and blood sample volume and quality of a refined retro-orbital bleeding technique in rats using a lateral approach. Lab Anim (NY). 2014;43(2). doi:10.1038/laban.432

30. Sellers RS, Morton D, Michael B, Roome N, Johnson JK, Yano BL, et al. Society of Toxicologic Pathology position paper: organ weight recommendations for toxicology studies. Toxicologic pathology. 2007. doi:10.1080/01926230701595300

